# l-DOPA stimulates the dopaminergic phenotype in human retina

**DOI:** 10.1101/2020.10.14.339366

**Authors:** Bojana Radojevic, Margarita Mauro-Herrera, Lea D. Bennett

## Abstract

Retinal organoids derived from inducible pluripotent stem cells were used to gain insight into the role of l-DOPA during human retinal development. Dopaminergic gene expression was indicated by assessing two dopamine receptors (DRD1 and *DRD2*), DOPA decarboxylase (*DDC*), and tyrosine hydroxylase (*TH*) via quantitative reverse transcription-polymerase chain reaction at various developmental stages. TH transcript levels started to express around day (D) 42, reached maximal expression ∼D63 and then decreased thereafter. At D29, proliferating retinal progenitors expressed *DRD1, DRD2*, and *DDC* at various levels of mRNA throughout the day. In the presence of l-DOPA, D29 retinal organoids expressed *DRD1* but *DRD2* mRNA expression was suppressed. Additionally, l-DOPA upregulated *TH* mRNA prior to dopaminergic amacrine cell (DAC) development. After the appearance of DACs, l-DOPA phase shifted expression of *DRD2* and synchronized mRNA expression of *DDC, DRD2*, and *TH*. The present results suggest unique mechanisms for DA signaling at different stages of development in the human retina.

## Introduction

Dopamine (DA) is a critical neurotransmitter that integrates signaling in the retina to mediate physiological functions such as circadian entrainment, neurogenesis, photoreceptor disk shedding, intraocular pressure, and refractive development.^1-4^ DA acts through receptors to activate or inhibit distinct pathways thereby mitigating various functions during neurogenesis. In this way, neuronal excitability occurs by second messenger cascades coupled to DA receptors (DRs). DRs are g-protein coupled receptors (GPCRs), classified by the g-protein that they couple to: D1-”like” couple to Gαs and D2-like couple to Gαi. ^5-7^ Additionally, DA is a zeitgeber, or time setter that regulates the expression of circadian rhythm genes and is critically involved in mediating the core molecular clock throughout the body. In adults, disruption of the DA pathway is associated with visual deficits in Parkinsonians disorders and schizophrenia.^8^ During embryogenesis, dysfunction in DA signaling has been linked to myopia,^4^ and ocular albinism^9^.

Research has revealed various processes in neurogenesis that are linked to DA signaling but these events depend on cell type and stage of development. For instance, DA increases neuronal proliferation when D2-like receptors are activated, but D1-like receptor activation decreases proliferation.^10^ Neurons that are GABAergic will migrate when DA binds D1-like receptors but this migration is inhibited by a D2-like receptor agonist.^11^ In cerebral cortical neurons, DA promotes axon growth by D2-like receptor activation which is inhibited with D1-like receptor stimulation.^12,13^ In striatal neurons, the opposite occurs; neurite outgrowth is stimulated by DA binding to D1-like receptors, but not with D2-like receptor activation.^14^ The basis for these effects in neural tissues of rodents is not fully understood and even less is known about DA mechanisms within developing human retinal neurons. The changing roles of this neuromodulator during development may be a fine-tuning mechanism to ensure cell migration and correct placement within the retina to achieve lamination.

DA synthesis occurs in a 2 step reaction whereby tyrosine hydroxylase (TH) converts tyrosine to l-DOPA. Next, DOPA decarboxylase (DDC) acts on l-DOPA to form DA (**Fig. 1**). DA is released and binds to its receptors on the post-synaptic neuron (**Fig. 1**). In the retina, seminal experiments using chicks have determined that early in embryogenesis extrinsic l-DOPA is made and secreted from the retinal pigmented epithelium (RPE).^15^ This upregulates DA synthesis and occurs prior to the appearance of dopaminergic amacrine cells (DACs). Additionally, at later stages during chick embryogenesis, DA inhibits axonal growth cones^16^ and limits the total number of amacrine cells that can synthesize DA.^17^ Taken together, these studies suggest that DA has different molecular affects that depend on cell type, developmental stage, and the local environment.

**Figure 1.**
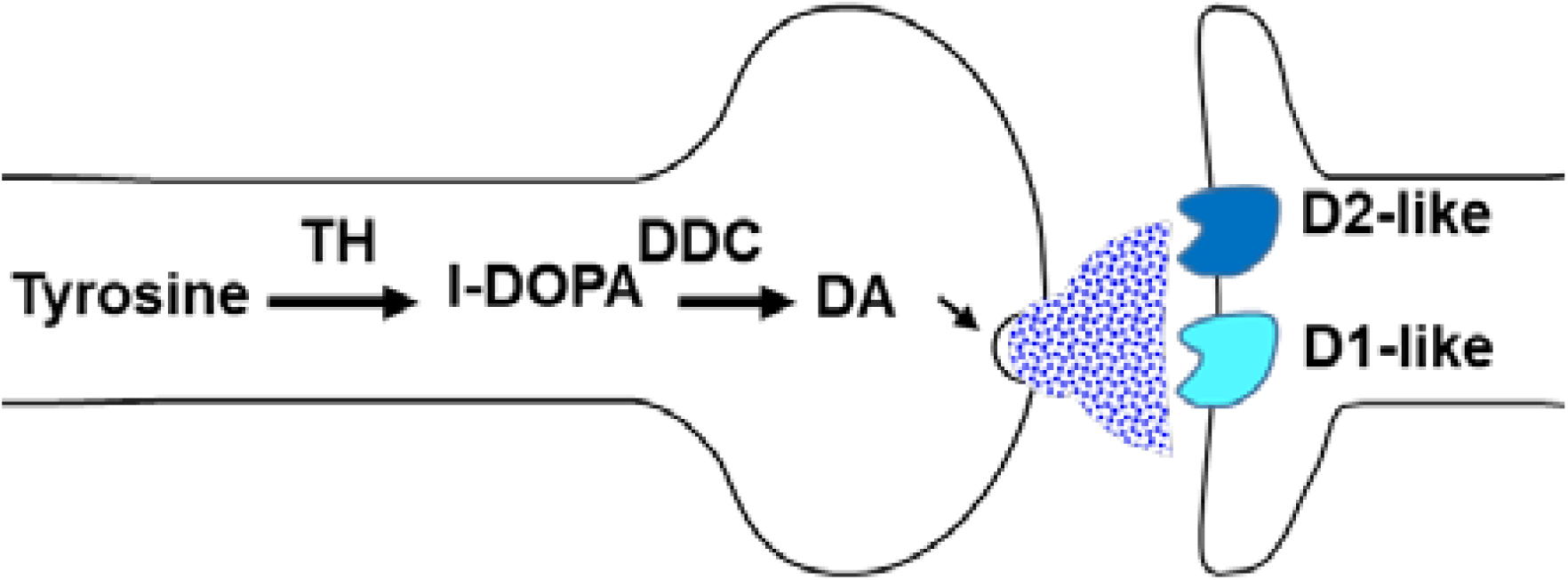
Dopamine biosynthesis.

In the present study, gene expression of *DRD1, DRD2, DDC*, and *TH* define the dopaminergic phenotype. Here we investigate the development of the dopaminergic phenotype and gain insight into the role of l-DOPA using human retinal tissue derived from human inducible pluripotent stem (iPS) cells. These results show that prior to the appearance of DACs, l-DOPA upregulates mRNA expression of *TH* and *DRD1* but suppresses *DRD2*. After the appearance of DACs, l-DOPA stimulates a phase shift so that *DRD2* is expressed maximally and synchronized with mRNA transcription of *DDC* and *TH*. Altogether, these results suggest unique mechanisms for DA signaling at different stages of development in the human retina.

## Methods

### Differentiation protocol

Human iPS cells were maintained in 6-well plates on Matrigel (Corning). We used two different iPS cell lines GM23270 and GM25256 (Coriel Institute). StemFlex (Thermo Fisher) media was used to maintain the health and pluripotency of the iPS cells and ReLeSR (STEMCELL Technologies) was used for passaging. Differentiation experiments were initiated using an established protocol.^18,19^ Briefly, embryoid body (EB) formation began with transition into neural induction medium [NIM; DMEM:F12 1:1, 1% N2 supplement, 1× MEM nonessential amino acids (MEM NEAA), 1× GlutaMAX (Thermo Fisher) and 2 mg/ml heparin (Sigma)] for 5 days. On day 6 (D6), 1.5 nM Bone Morphogenetic Protein (BMP4, R&D Systems) was added to fresh NIM. On d7, EBs were plated on Matrigel. Half of the media was replaced with fresh NIM on D9, d12, and d15. On D16, the media was changed to retinal differentiation medium (RDM;DMEM:F12 3:1, 2% B27 supplement, MEM NEAA, 1× antibiotic, antimycotic (Thermo Fisher) and 1× GlutaMAX). Retinal organoids that display an outer rim of neural retina were identified morphologically by light microscopy and dissected with a MSP ophthalmic surgical knife (Surgical Specialties Corporation) between D25-D30. Organoids were maintained in poly-HEMA-coated flasks (polyHEMA from Sigma) with twice-weekly feeding of 3D-RDM (DMEM:F12 3:1, 2% B27 supplement, 1× MEM NEAA, 1× antibiotic, anti-mycotic, and 1× GlutaMAX with 5% FBS, 100 μM taurine, 1:1000 chemically defined lipid supplement (11905031, Thermo Fisher)). All-trans retinoic acid (1 μM; Sigma) was included in the media until D120.

### Quantitative real-time polymerase chain reaction (RT-qPCR)

Gene expression was assessed by quantitative RT-qPCR. Retinal organoids (n=3-4) for each time point were homogenized using a Dounce Tissue Grinder (Sigma-Aldrich, UK) and processed using SYBR™ Green Fast Advanced Cells-to-CT ™ Kit (Invitrogen) to make cDNA. RT-qPCR was performed in triplicate using a CFX96 Real-Time System (Bio-Rad). Each primer (Table 1) was used at a final concentration of 1 µM. The reaction parameters were as follows: 50°C for 2 minutes, 95°C for 10 minutes to denature the cDNA and primers, 40 cycles of 95°C for 3 seconds followed by primer specific annealing temperature for 30 seconds (60°C), succeeded by a melt curve. A comparative cycle threshold (Ct)^20^ method was used to calculate the levels of expression that were normalized to *GAPDH* and relative to 0.

**Table 1.**
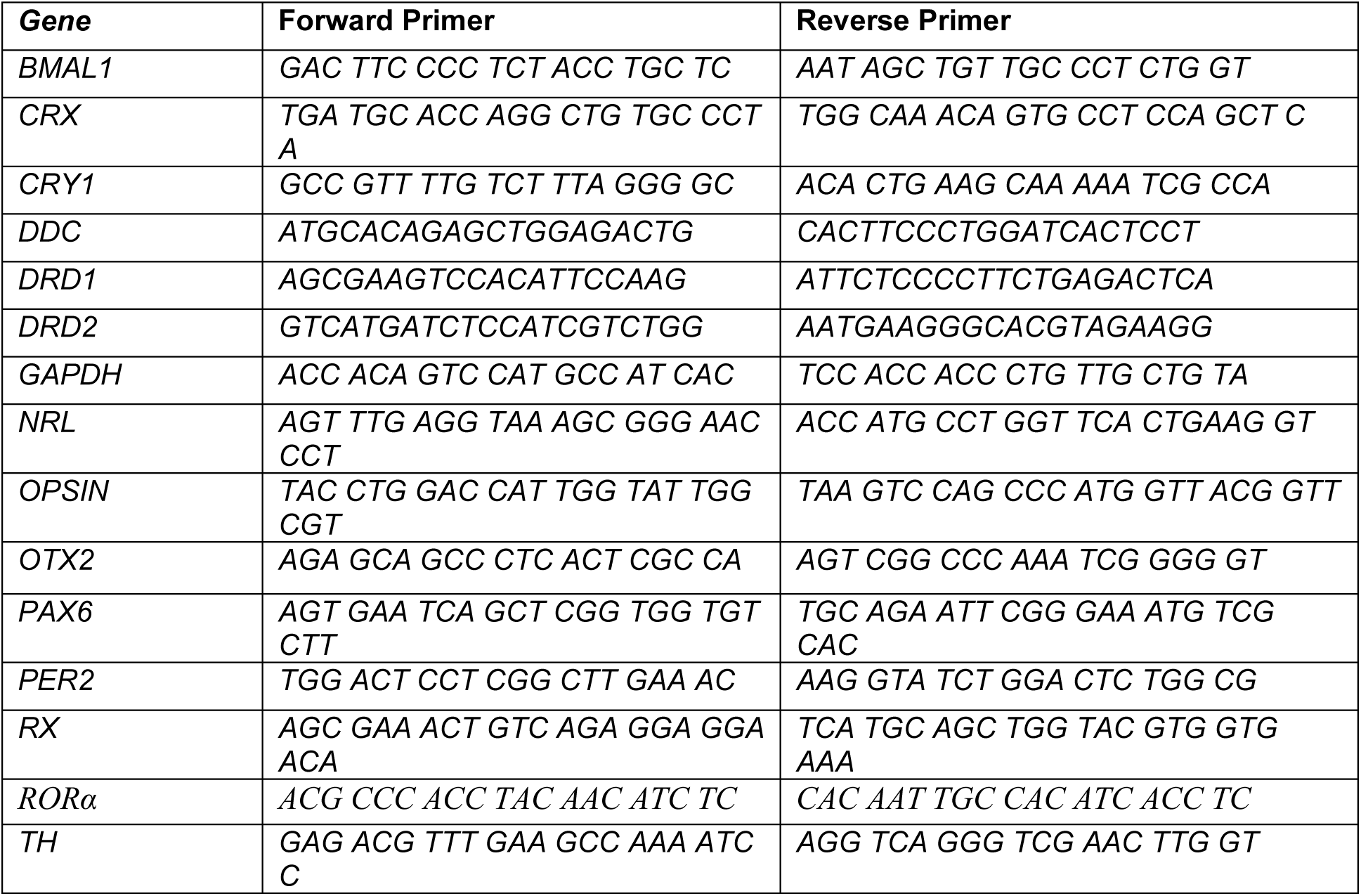

### Immunohistochemistry (IHC)

Retinal organoids were fixed in 4% paraformaldehyde (FD neuroTechnologies) at room temperature (RT) with gentle agitation for 35-60 minutes and washed 3× with PBS. Subsequently, retinal organoids were incubated in 15% sucrose in PBS for 1-2 hours, transferred to 30% sucrose, and stored at 4°C overnight. Retinal organoids were embedded in optimal cutting temperature compound and frozen at −20°C. Ten-micrometre cryostat sections were collected using a Leica cryostat onto Superfrost Plus slides and stored at − 20°C in slide boxes prior to immunostaining. Cryosections were air-dried, washed several times in PBS and incubated in blocking solution (10% normal donkey serum (NDS), 5% bovine serum albumin, 1% fish gelatin and 0.5% Triton X-100) for 1-2 hours at RT. Primary antibodies were incubated at 4°C overnight. See Table 2 for a list of primary antibodies, sources and concentrations. Secondary antibodies were diluted to 1:500 and added to tissues for 30 minutes in the dark at RT (Alexa Fluor 488, AF546 and AF647; Thermo Fisher) and again washed with PBS (3 × 10 minutes). Samples were incubated in DAPI (1:1000, Thermo Fisher Scientific) for 5 minutes, and then washed with PBS (3 × 10 minutes). Cover slips were mounted over the glass slides, then dried at RT and stored at 4 °C for microscopic observation. Samples were imaged on an Olympus FV1200 confocal microscope.

**Table 2.**

| **Antibodies** | **type** | **source** | **Cat number** | **Dilution** |
| --- | --- | --- | --- | --- |
| BRN3 | Rabbit polyclonal | Thermofisher | PA541509 | 1:50 |
| CaR | Rabbit polyclonal | Milipore | AB5054 | 1:500 |
| CRX | Mouse monoclonal | Abnova | H00001406-M02 | 1:100 |
| Ki-67 | Mouse monoclonal | BD Biosciences | 556003 | 1:500 |
| LHX2 | Mouse monoclonal | Santa Cruz | sc-81311 | 1:50 |
| OTX2 | Goat polyclonal | R&D | AF1979 | 1:2000 |
| PAX6 | Mouse monoclonal | DSHB | Pax6-s | 1:50 |
| SNCG | Mouse monoclonal | Abnova | H00006623-M01 | 1:500 |

### l-DOPA treatment

l-DOPA was added to the media (1mM, Sigma Aldrich) with retinal organoids on D29 and removed after 12h by changing the media. 3-5 retinal organoids were collected at different time points and assessed by RT-qPCR described above. To avoid neurotoxicity from excess l-DOPA or DA, we therefore administered l-DOPA at lower concentration (500µM) and reduced incubation time (4 hours) for D62 retinal organoids; retinal organoids were collected every 6hours during 48hours and processed as mentioned above.

## Results

To determine the appearance of dopaminergic cells, *TH* (rate-limiting enzyme in DA biosynthesis^21,22^) mRNA was quantified on different developmental days. Retinal organoids were harvested at the same clock time (CT; 8am) each day. *TH* started to develop ∼D42, was greatest ∼D63 and decreased by ∼D70 (**Fig. 2A**). At the onset of TH expression (D42), these retinal organoids were comprised of proliferating (*LHX2, Ki67*) neural progenitors (*PAX6*) and cells committed to the retinal ganglion cells (RGC) lineage (*BRN3*, **Fig. 2B**).^23^

**Figure 2.**
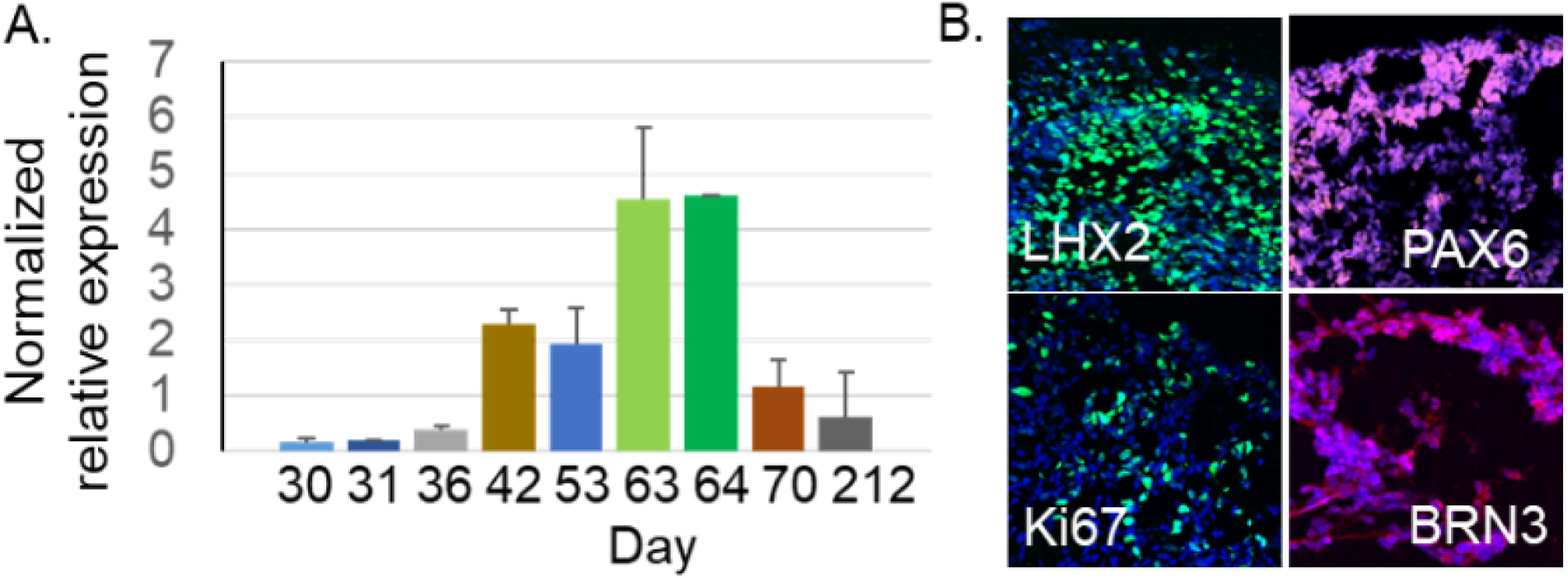
**A)** D30–D212 *TH* mRNA ± SD normalized (GAPDH); triplicates. n=3-4 **B)** IHC of proliferating (LHX2, PAX6, Ki67) and RGC-committed cells (BRN3).

Because DA is directly involved in regulating expression of circadian rhythm gene expression, we wanted to determine whether core clock genes (*BMAL1, PER2, CRY1*, and *RORα*) were rhythmically transcribed before the birth of dopaminergic cells. Therefore, these genes were assessed over a 24-hour period when retinal organoids were at D29 starting at 0 hour. At hours 4 and 8, *BMAL1* and *CRY1* showed lower mRNA levels than *PER2* and *RORα* (**Fig. 3A**). This trend reversed at hours 16 and 24. Additionally, dopaminergic gene expression of *DRD1, DRD2, DDC*, and *TH* was also evaluated in these D29 retinal organoids. *TH* was minimally detected but transcription for *DDC* and *DRD1* was upregulated at hours 4 and 12 (**Fig. 3B**). *DDC* and *DRD1* mRNA levels decreased at hour 16 when *DRD2* transcription was upregulated. Next, we wanted to determine if the expression of these genes would change when D29 retinal organoids were exposed to a zeitgeber. In other words, we wanted to know if these developing retinal cells were competent for circadian entrainment. Therefore, beginning at zeitgeber time 0 (ZT0), we added 1mM l-DOPA to the retinal organoids. Transcription of the core clock genes was diminished in the presence of l-DOPA at ZT4 (**Fig. 3C**). However, at ZT12, *CRY1* transcription was upregulated while the other genes continued to express at low levels (**Fig. 3C**). Dopaminergic gene expression was also modified by l-DOPA. *DRD1* and *DDC* continued to be expressed in the presence of l-DOPA at ZT4 (orange and blue lines, respectively; **Fig. 3D**). Conversely, expression of *DRD2* (gray line) was repressed by l-DOPA (**Fig. 3B, D**). Moreover, l-DOPA stimulated expression of *TH* at ZT12 (**Fig. 3D**, yellow line). D29-D30 retinal organoids expressed mRNA that confirmed neural induction (*PAX6*), eye field specification (*RX*), photoreceptor lineage (*CRX*), and anterior neural specification (*OTX2*). There was an absence of mature cone photoreceptors (*OPSIN*) and rod precursors (*NRL*; **Fig. 3E**). Therefore, the retinal organoids were highly proliferative and unlikely to contain cells that had terminally differentiated at D29.

**Figure 3.**
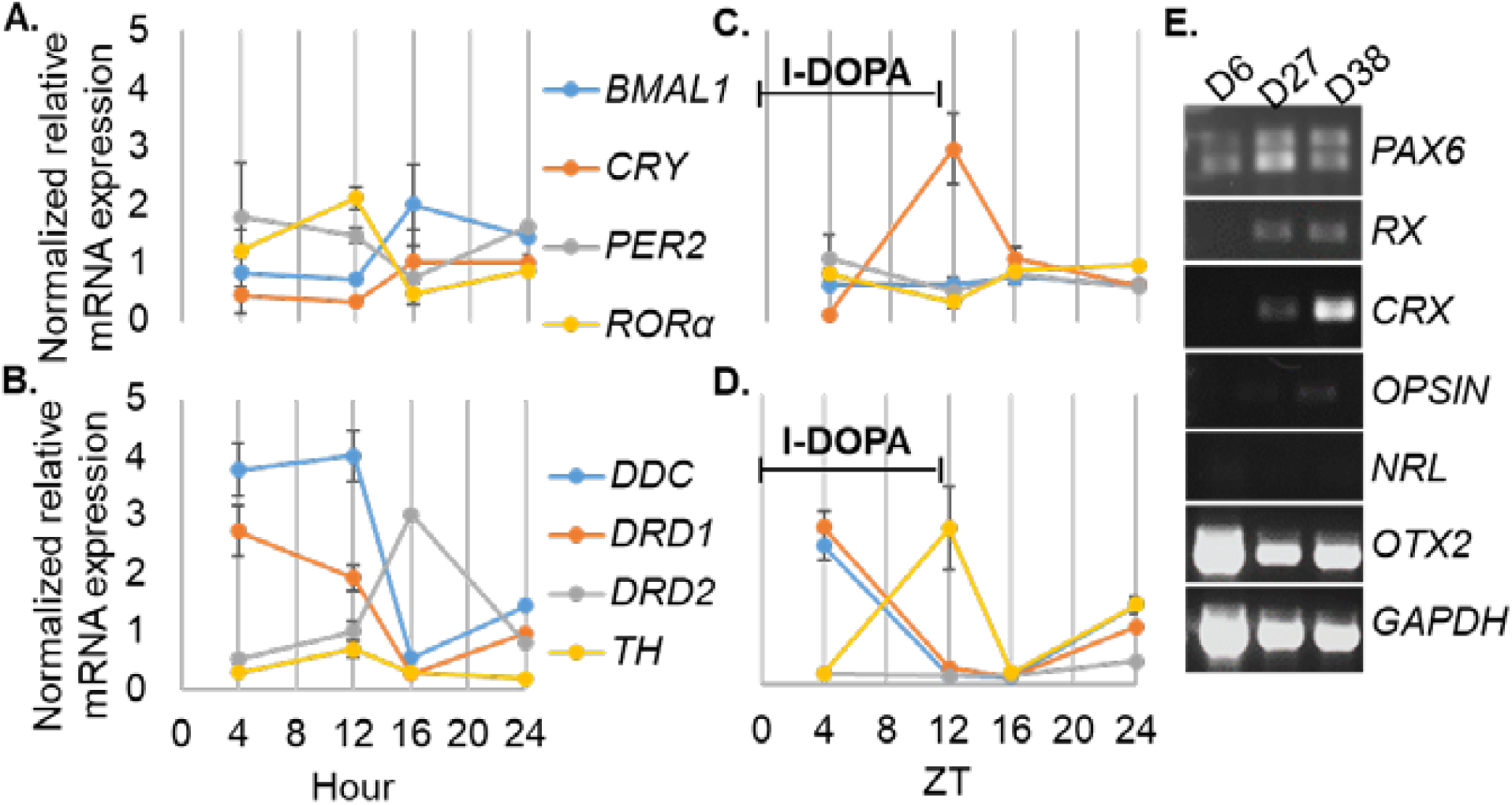
**A)** D29 retinal organoids mRNA expression of core clock and **B)** dopaminergic genes **C)** l-DOPA changed the pattern of gene expression of clock genes and **D)** downregulated *DRD2*. ZT, zeitgeber time (hours) **E)** RT-PCR showing neural progenitors.

Next, we wanted to determine if retinal organoids rhythmically transcribed dopaminergic genes after the birth of dopaminergic cells, indicated by *TH* expression. Therefore, at D64, retinal organoids were analyzed over 48 hours for expression of *DRD1, DRD2, DDC*, and *TH*. At CT24 and CT48, *TH* was upregulated (**Fig. 4A**). Expression of *DRD2* increased at CT30 (**Fig. 4A**). *DDC* was transcribed at the same times (CT24, CT30, and CT48). Assessment of *BMAL1* and *RORα* revealed maximal expression at CT 24 and CT30, respectively (**Fig. 4B**).

**Figure 4.**
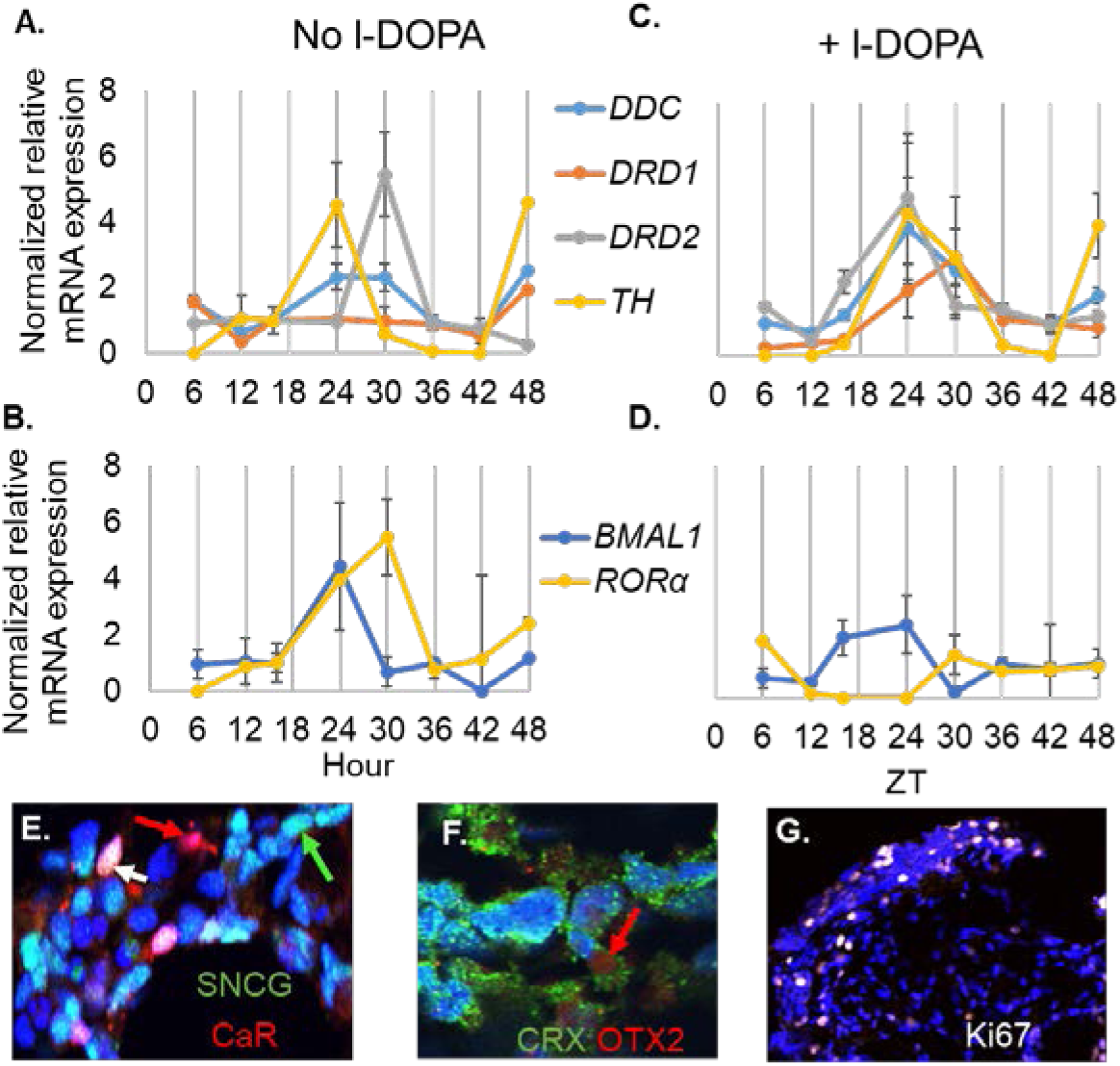
D62 retinal organoids. mRNA expression **A), B), C)** and **D)** 4 hours incubation l-DOPA, ZT, zeitgeber time (hours). **E)** IHC D62 showing SNCG (RGCs, green arrow), CaR (RGC, white arrow; amacrine cell, red arrow). **F)** Photoreceptor progenitors CRX and OTX2 (red arrow). **G)** Minimal mitotic proliferation (Ki67)

To evaluate the effect of l-DOPA (500µM for 4 hours) on retinal organoids at D64, we quantified gene expression of the dopaminergic phenotype. mRNA expression of *DRD2, DDC*, and *TH* were synchronized to express maximally at ZT24 whereas *DRD1* showed a delay in expression that peaked at ZT30 (**Fig. 4C**). Expression of *BMAL1* was antiphase to *RORα* after l-DOPA treatment (**Fig. 4D**). IHC at D62 indicated the presence of RGCs (SNCG^+^/CaR^-^, green arrow and SNCG^+^/CaR^+^, white arrow), amacrine cells (SNCG^-^/CaR^+^, red arrow; **Fig. 4E**) and photoreceptor progenitors (CRX and OTX2, red arrow; **Fig. 4F**). Differentiated cells at this developmental stage was also supported by minimal detection mitotic proliferation (Ki67; **Fig. 4G**). Therefore, D64 retinal organoids consisted mainly of terminally differentiated inner retinal cells.

## Discussion

Here we show that DA signaling genes (*DRD1, DRD2, TH, DDC*) begin to express autonomously around D42-D70, without extrinsic l-DOPA. Furthermore, in the proliferating retinal cells, extrinsic l-DOPA triggered mRNA expression of DA signaling genes. This has also been reported in chick retinae. Kubrusly *et al*^15^ showed that the retinal pigmented epithelium (RPE) supplies l-DOPA prior to the birth of dopaminergic cells which stimulates DA synthesis.^15^ However, whether l-DOPA is released from human RPE during development has yet to be determined.

It has been suggested that the clock could play a major role in determining the timing of cell differentiation during development.^24^ Recently, core clock proteins have been postulated to restrict oscillations in undifferentiated cells^25^ but upon differentiation, may guide neuron migration and synaptic network formation.^26^ In *vivo* and in synchronized cells, *PER1/2* and *CRY1/2* are upregulated during the subjective day so that they can form a complex that subsequently negatively regulate the expression of clock genes.^27-29^ We did not find parallel expression of *CRY1* and *PER2* at D29 and l-DOPA did not synchronize mRNA expression of these genes (**Fig. 3**). Additionally, the retinoic acid-related orphan receptor (RORα, β, c) protein is known to stimulate gene expression of *BMAL1* which is occurs during the subjective night.^3,30-32^ However, we did not find *BMAL1* transcription to be increased following the upregulation of *RORα* mRNA expression at D29. Again, l-DOPA did not stimulate *RORα* mRNA expression nor subsequent *BMAL1* transcription at D29. Conversely, *CRY1* expression was upregulated by l-DOPA. It has been suggested that during development, CRY1 has regulatory functions that are independent of the circadian transcription/translation feedback loop that is active in almost every cell in adults.^33^ As such, l-DOPA may be required for CRY1 expression during embryogenesis. The D29 retinal organoids were comprised of undifferentiated and proliferating progenitor cells. Altogether, these results support the notion that the role of circadian mechanisms in early retinogenesis (before cells terminally differentiate) is to restrict oscillations and prevent synchronization of circadian gene expression.

At D29, the retinal organoids were highly proliferative and at a stage of development prior to inner retinal cell differentiation. We do not know if expression of *TH*, albeit was extremely low, had functional significance. To determine whether DA is synthesized and secreted from proliferating progenitors as well as from DACs, we need to evaluate DA (mass spectrometry) in media and retinal organoids at ∼D29 and ∼D64, respectively. Nevertheless, even with the low levels of *TH*, D29 retinal organoids expressed *DDC, DRD1* and *DRD2*. Specifically, expression of *DRD1* mRNA was higher when *DRD2* showed lower expression. Conversely, when transcription of *DRD1* was decreased, *DRD2* mRNA increased in D29 retinal organoids (**Fig. 3**). On the other hand, D62 retinal organoids showed a preference for *DRD2* mRNA expression and not *DRD1* (**Fig. 4A** and **3C**). Moreover, D62 retinal organoids responded differently to l-DOPA treatment compared to D29. At this stage, l-DOPA did not directly stimulate the expression of *DRD1*, but instead, acted as a zeitgeber by phase shifting the expression of *DRD2, BMAL1*, and *RORα* (**Fig. 4**). Unlike D29, at D64, transcription of *BMAL1* and *RORα* was set by l-DOPA where increased *RORα* mRNA expression occurred when *BMAL1* was downregulated. Conversely, when *BMAL1* increased mRNA expression, *RORα* was downregulated (**Fig. 4D**). It is uncertain whether a single treatment with l-DOPA will be sufficient to synchronize cyclic dopaminergic gene expression in retinal organoids. In order to establish rhythmicity, these measures need to be assessed for at least 5 days including time points throughout each day.^34^ Altogether, results from this study show unique DA signaling at different stages of development in the human retina. This suggests that the specific pathways triggered by DA mediates distinct physiological processes during retinal neurogenesis that varies among proliferating, differentiating, and terminally differentiated retinal neurons.

